# Intention to learn differentially affects subprocesses of procedural learning and consolidation: Evidence from a probabilistic sequence learning task

**DOI:** 10.1101/433243

**Authors:** Kata Horváth, Csenge Török, Orsolya Pesthy, Dezso Nemeth, Karolina Janacsek

**Affiliations:** Doctoral School of Psychology, ELTE Eötvös Loránd University, Izabella utca 46, H–1064, Budapest, Hungary; Institute of Psychology, ELTE Eötvös Loránd University, Izabella utca 46, H–1064, Budapest, Hungary; Brain, Memory and Language Research Group, Institute of Cognitive Neuroscience and Psychology, Research Centre for Natural Sciences, Hungarian Academy of Sciences, Magyar tudósok körútja 2, H–1117, Budapest, Hungary; Lyon Neuroscience Research Center, INSERM, CNRS, Université de Lyon, Centre Hospitalier Le Vinatier - Bâtiment 462 - Neurocampus 95 boulevard Pinel 69675 Bron, Lyon, France

**Keywords:** intention to learn, incidental learning, sequence learning, statistical learning, consolidation, sleep

## Abstract

Procedural memory facilitates the efficient processing of complex environmental stimuli and contributes to the acquisition of automatic behaviours and habits. Learning can occur intentionally or incidentally, yet, how the mode of learning affects procedural memory is still poorly understood. Importantly, procedural memory is a complex cognitive function composed of different subprocesses, including the acquisition and consolidation of *statistical, frequency-based* and *sequential, order-based knowledge*. Therefore, we tested how statistical and sequence knowledge develops during incidental versus intentional procedural memory formation and during consolidation. Seventy-four young adults performed either the uncued, incidental (N = 37) or the cued, intentional (N = 37) version of a probabilistic sequence learning task. Performance was retested after a 12-hour offline period, enabling us to test the effect of sleep on consolidation; therefore, half of the participants slept during the delay, while the other half had normal daily activity (PM-AM versus AM-PM design). The mode of learning (incidental versus intentional) had no effect on the acquisition of statistical knowledge, while intention to learn increased sequence learning performance. Consolidation was not affected by intention to learn: Both statistical and sequence knowledge was retained over the 12-hour delay, irrespective of the mode of learning and whether the delay included sleep or wake activity. These results suggest a time-dependent instead of sleep-dependent consolidation of both statistical and sequence knowledge. Our findings could contribute to a better understanding of how the mode of learning (intentional or incidental) affects procedural memory formation and consolidation.

## Introduction

Procedural memory underlies the acquisition of and adaptation to the complex regularities in our environment. Learning in the procedural memory system can take place incidentally (i.e., without intention to learn and/or awareness that learning occurs) or intentionally (Fletcher et al., 2005; D. Howard & Howard, 2001). It is still debated whether incidental and intentional learning constitute two distinct processes of procedural memory (Reber & Squire, 1994, 1998) or the acquired knowledge taps into the same procedural memory system, irrespective of whether learning occurred incidentally or intentionally (Cleeremans, 2006; Henke, 2010; Perruchet & Pacton, 2006). To date, only a few studies focused on the direct investigation of this question and led to mixed findings (Ferdinand, Rünger, Frensch, & Mecklinger, 2010; Robertson, Pascual-Leone, & Press, 2004; Rüsseler & Rösler, 2000; Sanchez & Reber, 2013). Focusing on incidental versus intentional learning, our main goal was to clarify these mixed results, and examine *how the mode of learning affects procedural memory* in a unified, controlled paradigm that goes beyond previous study designs in two ways.

First, the concept of procedural memory is an umbrella term covering several subprocesses that could be distinguished both on conceptual and methodological level (Kóbor et al., 2018; Nemeth, Janacsek, & Fiser, 2013; Simor et al., 2019). The contradictory findings often observed in procedural memory research may at least partly result from the lack of such differentiation. Here we aimed to differentiate between at least two subprocesses of procedural memory, namely *the acquisition of statistical and sequence knowledge* (Nemeth, Janacsek, & Fiser, 2013), and assess these processes in parallel. Statistical learning refers to the recognition and acquisition of frequency or probability based shorter-range associations among stimuli (e.g., differentiating between more frequent and less frequent stimulus chunks, such as pairs or triplets of stimuli; Armstrong, Frost, & Christiansen, 2017; Fiser & Aslin, 2002; Thiessen, Kronstein, & Hufnagle, 2013; Turk-Browne, Scholl, Johnson, & Chun, 2010), while sequence learning refers to the acquisition of a series of repeating elements occurring in the same order without or with some embedded noise (deterministic vs. probabilistic sequences, respectively; Brawn, Fenn, Nusbaum, & Margoliash, 2010; Rickard, Cai, Rieth, Jones, & Ard, 2008). Recent findings indicate that statistical and sequence learning follow different developmental trajectories (Nemeth, Janacsek, & Fiser, 2013) and show different electrophysiological patterns (Kóbor et al., 2018; Simor et al., 2019). In this study, we used a probabilistic sequence learning task, which is a suitable tool to measure both statistical learning and sequence learning within procedural memory, due to the embedded alternating regularity.

To the best of our knowledge, only one study tested the effect of intention to learn on the acquisition of statistical and sequence knowledge in parallel, using the same task, and found that intentional learning improved both knowledge compared to incidental learning in healthy young adults (Nemeth, Janacsek, & Fiser, 2013). In this study, however, participants had as much time as they needed for stimulus processing and responding, which may have favoured intentional learning processes over incidental learning, and potentially amplified the group differences. To overcome these limitations, here we aimed to test whether the mode of learning (incidental vs. intentional) differentially affects the two subprocesses of procedural memory (statistical and sequence learning) while controlling for the time on task across the incidental and intentional learning conditions.

Second, beyond focusing on how the mode of learning affects the acquisition of statistical and sequence knowledge, we also aimed to test its effect on the *consolidation* of these subprocesses. Consolidation refers to the changes in the acquired knowledge during the post-learning offline periods (Krakauer & Shadmehr, 2006; Meier & Cock, 2014; Robertson, 2009; Robertson, Pascual-Leone, & Miall, 2004). Typically, incidentally acquired (statistical or sequence) knowledge is retained during the offline period (Hallgató, Győri-Dani, Pekár, Janacsek, & Nemeth, 2013; Janacsek & Nemeth, 2012; Kim, Seitz, Feenstra, & Shams, 2009; Kóbor et al., 2019; Nemeth & Janacsek, 2010; Nemeth et al., 2010; Rickard et al., 2008; Song, Howard, & Howard, 2007b). In contrast, previous studies on intentionally acquired sequence knowledge led to mixed findings: while in some cases successful consolidation of the acquired knowledge means lower degree of forgetting (Mednick, Cai, Kanady, & Drummond, 2008), in other cases retention or even improvement in performance can occur after the delay period (Pan & Rickard, 2015; Robertson, Pascual-Leone, & Miall, 2004). To the best of our knowledge, consolidation of statistical knowledge acquired under intentional learning condition has been investigated solely after a short-term delay period (i.e. 1.5 hour), where retention was observed (Simor et al., 2019).

Moreover, the activity during the delay period may also alter the consolidation of procedural knowledge: although a great body of research showed that sleep does not affect procedural memory on behavioural level (Csábi et al., 2016; Hallgató et al., 2013; Nemeth et al., 2010; Peigneux et al., 2003, 2006; Simor et al., 2019; Song et al., 2007b), other studies argued for a beneficial effect of sleep in this process, particularly if learning occurred intentionally (Durrant, Taylor, Cairney, & Lewis, 2011; King, Hoedlmoser, Hirschauer, Dolfen, & Albouy, 2017; Robertson, Pascual-Leone, & Press, 2004; Spencer, Sunm, & Ivry, 2006). Overall, it appears that offline improvement is more often observed for sequence learning than for statistical learning. Furthermore, sequential regularities may easily become explicitly recognized (consciously accessible), which seems to consolidate in a more effective manner during sleep compared to an awake delay period.

In a recent study, using only the intentional learning condition, Simor et al. (2019) compared statistical and sequence learning in the same task and tested their consolidation by retesting performance after a short period containing either napping, quiet rest or active awake activity. They reported behaviourally similar performance after the delay period in all three groups. Importantly, consolidation of intentionally vs. incidentally acquired knowledge was not contrasted in that study. Moreover, napping usually does not contain REM sleep due to its short length, possibly resulting in different consolidation effects as opposed to the effects of a full night sleep (Tucker et al., 2006). To systematically test these possibilities, we examined whether the intention to learn differentially affects the consolidation of statistical versus sequence knowledge, taking into account the activity during the consolidation period (sleep vs. wake) as well.

Overall, the aim of the present study was thus twofold. First, we aimed to investigate whether the mode of learning (intentional vs. incidental) differentially affects statistical learning and sequence learning while controlling for the time on task. Second, we aimed to test not only how statistical and sequence knowledge emerges during the learning phase but also how the acquired knowledge consolidates over a delay period while taking into account the possible role of sleep. We employed a widely used probabilistic learning task, namely the Alternating Serial Reaction Time (ASRT) task, to assess these processes. To provide a fuller picture of how the intention to learn and sleep vs. wake offline periods affect the acquisition and consolidation of statistical and sequence knowledge, we also examined whether the acquired statistical and sequence knowledge has become consciously available using additional tasks beyond the classical RT and accuracy measures of the ASRT task. We hypothesized that intention to learn helps the acquisition of both statistical and sequence information, based on the findings of Nemeth et al. (2013). We also expected that (both statistical and sequence) knowledge acquired in the incidental learning condition is retained during the delay, irrespective of the delay activity (sleep vs. wake). Similarly, we expected that statistical knowledge acquired in the intentional learning condition is also retained regardless of whether the offline delay contained sleep or an awake period. In contrast, we hypothesized that the consolidation of sequence knowledge acquired in the intentional learning condition is sleep-dependent, with better consolidation after the sleep period compared to the awake delay period.

## Methods

### Participants

Ninety-three healthy young adults participated in the experiment. We matched the Incidental and Intentional groups based on age, education, gender, handedness, working memory and attention performance. Six participants were excluded due to matching (differences in age and working memory capacity) and another thirteen participants due to outlier performance on the ASRT task (average accuracy and/or raw reaction time separately for each trial type fell outside three SDs in at least two epochs out of five in the Learning Phase). Thus, the final sample consisted of seventy-four adults, thirty-seven in each group (Table 1).

**Table 1.**
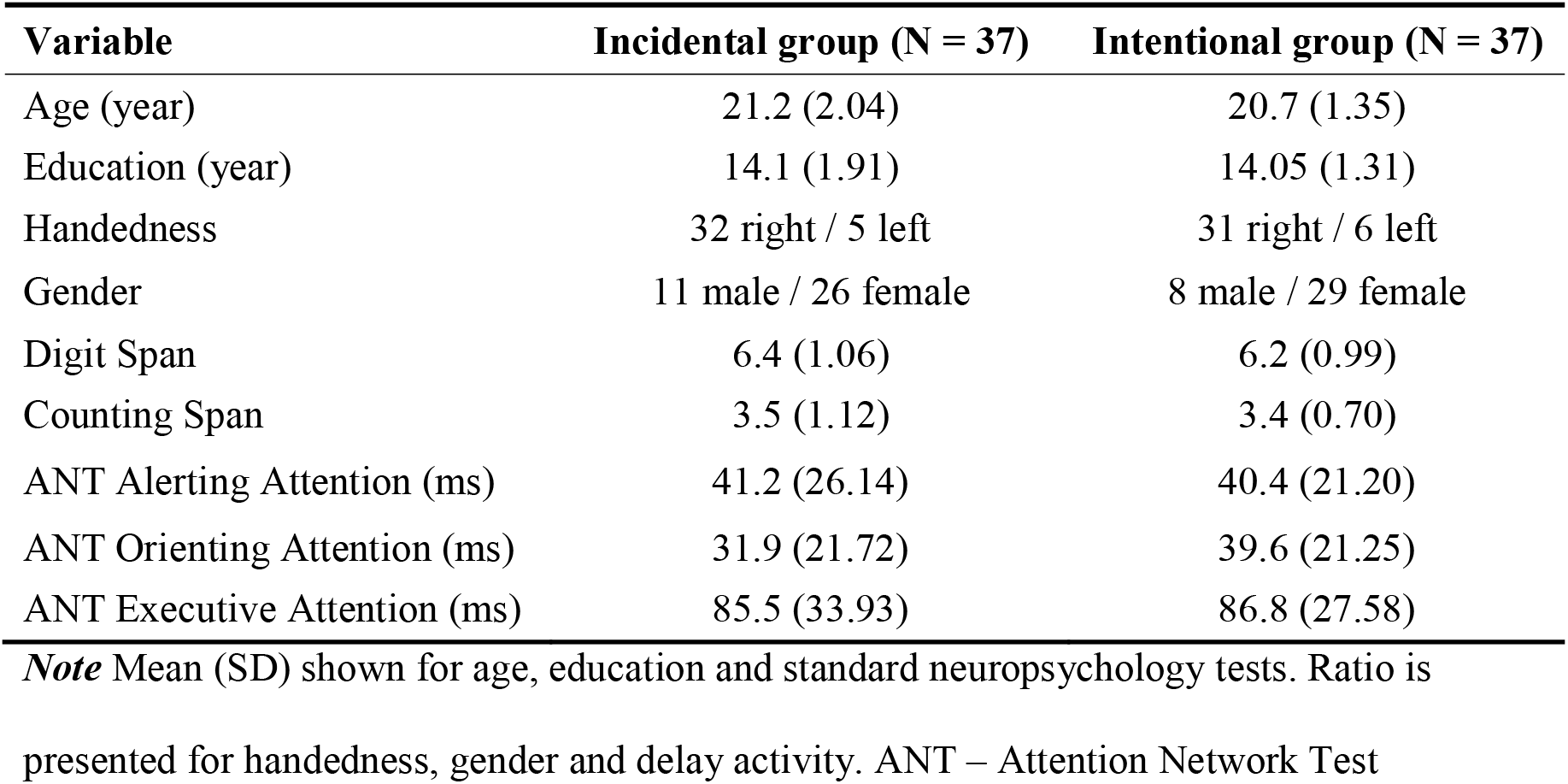
Demographic data of the Incidental and Intentional groups.

All participants had normal or corrected-to-normal vision, none of them reported a history of any neurological and/or psychiatric condition and drug-use. Prior to their inclusion in the study, participants provided informed consent to the procedure as approved by the research ethics committee of Eötvös Loránd University, Budapest, Hungary. The study was conducted in accordance with the Declaration of Helsinki and participants received course credits for taking part in the experiment.

Handedness was measured by the Edinburgh Handedness Inventory (Oldfield, 1971). General cognitive performance was measured by three neuropsychological tests: participants completed a digit span task (Isaacs & Vargha-Khadem, 1989; Racsmány, Lukács, Németh, & Pléh, 2005), a counting span task (Case, Kurland, & Goldberg, 1982; Conway et al., 2005; Engle, Tuholski, Laughlin, & Conway, 1999), and an attentional network test (Fan, McCandliss, Sommer, Raz, & Posner, 2002).

### Tasks

#### Alternating Serial Reaction Time (ASRT) task

The ASRT task was used to measure procedural memory. In this task, the target stimulus appeared in one of the four horizontally arranged circles on the screen. Participants were instructed to respond with the corresponding key (Z, C, B or M on a QWERTY keyboard) when the stimulus occurred. One block of the ASRT task contained 85 trials (stimuli). The presentation of the stimuli was determined by an eight-element sequence, in which pattern (P) and random (r) elements were alternating. In each block, the eight-element alternating sequence repeated 10 times after five warm-up trials consisting only of random stimuli. One example of the sequence is 2-r-3-r-1-r-4-r, where numbers represent predetermined locations on the screen and r indicate randomly chosen locations out of the four possible ones. This alternating structure resulted in some of three consecutive trials (*triplets*) occurring more frequently than others (*frequency property* of the task). If a triplet appeared frequently, its first two elements predicted the third element with a greater certainty (called *high-frequency triplets*, taking up 62.5% of all trials), while the third element of infrequent triplets was less predictable (called *low-frequency triplets*, taking up 37.5% of all trials). This frequency information defined the statistical property of the triplets. However, triplets could also be defined based on their structure: the last element of a triplet could occur in a pattern position (P-r-P structure) or in a random one (r-P-r structure; *sequential property* of the task). Based on the frequency and sequential properties of triplets, the last element of each triplet could be divided into three categories: pattern-high, random-high and random-low elements (Figure 1A).

**Figure 1.**
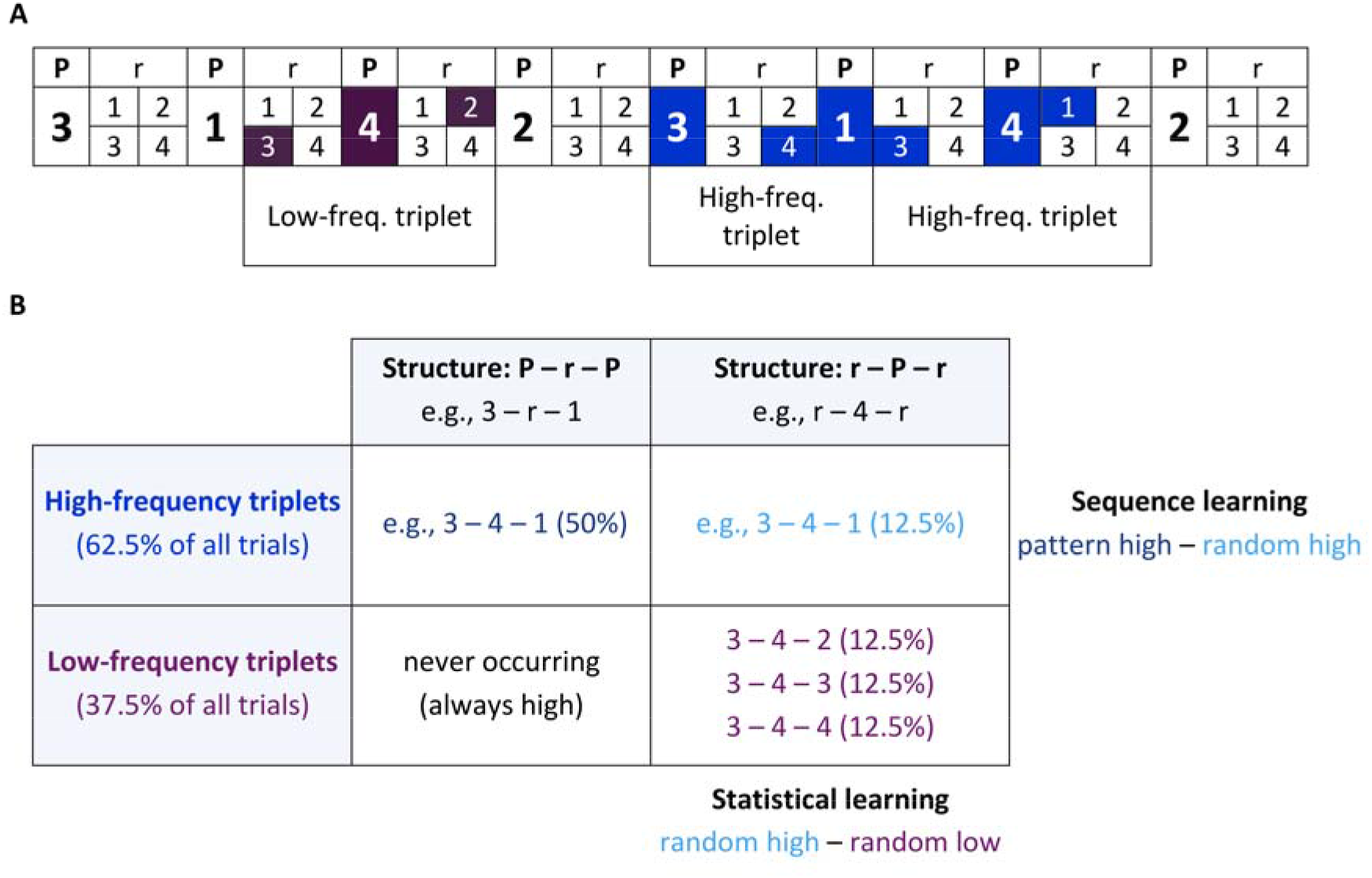
Stimulus structure and learning processes in the ASRT task. (P – pattern, r – random). **(A)** As the ASRT task contains an alternating sequence structure (e.g., 2r3r1r4r, where numbers correspond to the four locations on the screen and the r represents randomly chosen locations), some runs of three consecutive elements (called triplets) occur more frequently (blue) than others (purple). In the example above 2-_-4, 4-_-3, 3-_-1, and 1-_-2 occur often. In contrast, 2-_-3, 1-_-4 or 3-_-2 would occur less frequently. **(B)** We determined for each pattern and random stimulus whether it was the last element of a high- or a low-frequency triplet, thus three different elements could occur: pattern (dark blue, always high-frequency), random-high (light blue) and random-low (purple). Statistical learning is computed as the difference in responses between random-high and random low-elements. Sequence learning is defined as the difference in responses between pattern and random-high elements.

Different learning processes can be measured in the ASRT task. So far, studies mainly focused on *triplet learning*, however it is not a pure measure and at least two separate learning processes contribute to this learning measure, namely *statistical learning* and *sequence learning* (Kóbor et al., 2018; Nemeth, Janacsek, & Fiser, 2013; Simor et al., 2019). Statistical learning is defined as the difference in responses between high-frequency and low-frequency triplets analyzed only in random trial types. In this case, the sequential properties of the stimuli are controlled, and the only difference between the two stimulus types is frequency-based, that is purely statistical in nature. In contrast, sequence learning is defined as the differences in responses between pattern (always high-frequency) and high-frequency random trial types. Here, the statistical properties of the stimuli are controlled (both types are high-frequency), and the only differentiation of the two trial types is based on sequence properties (pattern vs. random; Figure 1B). Importantly, both learning measures – expressed as difference scores – can be separated from the so called *general skill learning* that is related to general performance improvement as the task progresses (mainly due to improved visuomotor coordination) and affects the different trial types similarly. Here we focus only on statistical learning and sequence learning.

It has been well documented that participants do not become aware of the underlying statistical/sequence structure embedded in the classic, incidental version of the ASRT task even after extended practice (e.g., ten days (Song, Howard, & Howard, 2007a)); and when examined with more sensitive recognition tests (D. Howard et al., 2004), thus it indeed measures incidental learning. Nevertheless, to ascertain that the sequence structure of the ASRT task and the learning situation itself remained fully implicit in the Incidental group in the current study, a short **questionnaire** was administered after the Testing Phase, similar to previous studies (Nemeth et al., 2010; Song et al., 2007a; Unoka et al., 2017). Participants were asked whether they observed any regularity in the task. None of them reported anything regarding the sequence embedded in the ASRT task, and moreover, none of them was aware of that any kind of learning occurred during the task.

To test whether the Intentional group successfully learned the sequence order, a **sequence report** was administered after each block of the ASRT task. We asked them to report the sequence of the pattern stimuli: they had to type the four-digit order of the sequential elements three times. However, participants were not informed about the length of the sequence, thus they had to figure out not only the order but the length of the sequence as well. We expected this sequence report to be more sensitive and avoid ceiling effects as seen in Nemeth et al. (2013) where information about the length was provided. Performance was defined as percentage of reporting the sequence correctly after a given block (12 items in correct order corresponding to 100% performance), and average performance was calculated for each epoch to facilitate comparison with the ASRT learning performance measures. Data from one participant was missing due to technical issues during data collection. Note that sequence reports after each block were not required in the Incidental group as those would have undermined the incidental nature of the task in this group.

#### Inclusion-Exclusion Task

Whereas sequence reports after each block in the Intentional group was administered to reveal how consciously accessible sequence knowledge (order information) became during learning, the Inclusion-Exclusion task (Destrebecqz & Cleeremans, 2001; Destrebecqz et al., 2005; Fu, Dienes, & Fu, 2010; Jiménez, Vaquero, & Lupiáñez, 2006) was administered to reveal how consciously accessible triplet knowledge (frequency-based information) became in both the Incidental and Intentional groups. This task is based on the well-established “Process Dissociation Procedure” (Jacoby, 1991) and consists of two parts: first, participants were asked to report in what order the stimuli (both pattern and random elements) appeared in the task (*Inclusion condition)*, then they had to generate a new sequence of stimuli (both pattern and random elements) that was different from the learned one (*Exclusion condition*). Both parts contained four runs and the participants used the same response buttons as in the ASRT task. Each run finished after 24 button presses, which was equal to three rounds of the eight-element alternating sequence. Since participants were asked to produce a stimulus stream including both pattern and random stimuli, this task enabled us to measure how consciously accessible their knowledge about the statistical regularities (i.e., triplets) was. Following the standard analysis and interpretation of the task outlined in previous studies, good performance in the Inclusion part can be achieved by solely implicit knowledge (i.e., without intentional access to their knowledge), while good performance in the Exclusion part requires accessible explicit knowledge to exert control over their responses and generate a stimulus stream that is indeed different from what they learned. Consequently, to test whether participants gained conscious knowledge of the statistical regularities, we calculated the percentage of high-frequency triplets in the Inclusion and Exclusion conditions separately, and tested whether participants produced more high-frequency triplets than it is expected by chance and whether the percentage of high-frequency triplets differed across (Inclusion/Exclusion) conditions or across groups (Kóbor, Janacsek, Takács, & Nemeth, 2017).

Data from five participants (two in the Incidental group and three in the Intentional group) was missing due to technical difficulties. Further eight participants (three in the Incidental group and five in the Intentional group) were excluded from analyses as they did not follow the instructions in the Exclusion condition.

### Procedure

The experiment consisted of two sessions to assess both learning and consolidation of the acquired knowledge. The Learning Phase contained 25 blocks. Half of the participants received the uncued version of ASRT (*Incidental group*), while the other half performed a cued version of the task (*Intentional group*). The Incidental group was informed that the aim of the experiment was to measure the effect of extended practice on motor performance, thus they were unaware of the sequential structure embedded in the task. In contrast, participants in the Intentional group were informed about the sequence structure of the task and were instructed to learn the sequence of the pattern elements. In this case, pattern and random stimuli were marked with different target pictures (dogs for pattern and penguins for random stimuli) similarly to the original study of Nemeth et al. (2013). We chose this design because a previous ASRT study showed that the intentional instruction alone does not lead to the discovery of the sequence order or can even disrupt learning performance (D. Howard & Howard, 2001). Moreover, based on unpublished data, the cued nature of the stimuli alone (i.e., without instruction to learn the sequence order) is not sufficient for intentional learning to occur either. Thus, the combined cuing and instruction appears to be necessary to ensure that participants in the intentional group can identify the pattern trials that follow a predetermined sequence but it does not warrant that participants can find and learn the sequential pattern itself (D. Howard & Howard, 2001; Song, 2009; Song et al., 2007a). Consequently, the Incidental and Intentional groups were named after the lack or presence of intention to learn, irrespective of whether or not they gained consciously accessible knowledge about the sequential regularities in the task.

We went beyond the previous studies by controlling the time on task with a fixed inter-stimulus interval (ISI) instead of the self-paced timing used previously since the latter enables the intentional group to spend more time on task. With this modification, we aimed to eliminate any potential confounds that could have affected the comparison of the Intentional vs. Incidental groups’ performance measures in previous studies. The timing of an experimental trial was the following: The duration of stimulus presentation was 500 ms (when participants were required to respond to the stimulus), then the four empty circles were presented for 120 m before the next stimulus appeared, thus, the total ISI was 700 ms. These values are defined based on previous studies investigating healthy young adults, where participants had an average response time under 450 ms at the beginning of the task and 430 ms by the end of the Learning Phase (Nemeth et al., 2010; Nemeth, Janacsek, Polner, & Kovacs, 2013; Tóth et al., 2017; Unoka et al., 2017).

The Learning Phase was followed by a 12-hour delay, thereafter the Testing Phase was administered, which contained five blocks of the ASRT task. In order to counterbalance potential time of day effects and measure the effect of sleep on consolidation processes, an AM-PM vs. PM-AM design was used: during the delay, half of both Incidental and Intentional groups slept (Sleep subgroups, PM-AM design) and the other half had normal daily activity (No-sleep subgroups, AM-PM design; Nemeth et al., 2010; Song et al., 2007b). Twenty participants formed the Incidental – No-sleep subgroup and 17 formed the Incidental – Sleep subgroup, while 19 participants formed the Intentional – No-sleep subgroup and 18 formed the Intentional – Sleep subgroup. All AM sessions took place between 7 AM and 10 AM, while all PM sessions took place between 7 PM and 10 PM. The four subgroups did not differ in age, education, gender, handedness, working memory or attention performance (all *p*s > .120). Participants in the sleep subgroups slept on average 6.0 hours (SD = 0.77). All participants were aware that they perform the same task in the second experimental session (Figure 2). Following the Testing Phase, the incidental group was asked whether they noticed any regularities in the task, then received information about the hidden sequence structure of the task. Then both the Incidental and the Intentional group performed the Inclusion-Exclusion Task.

**Figure 2.**
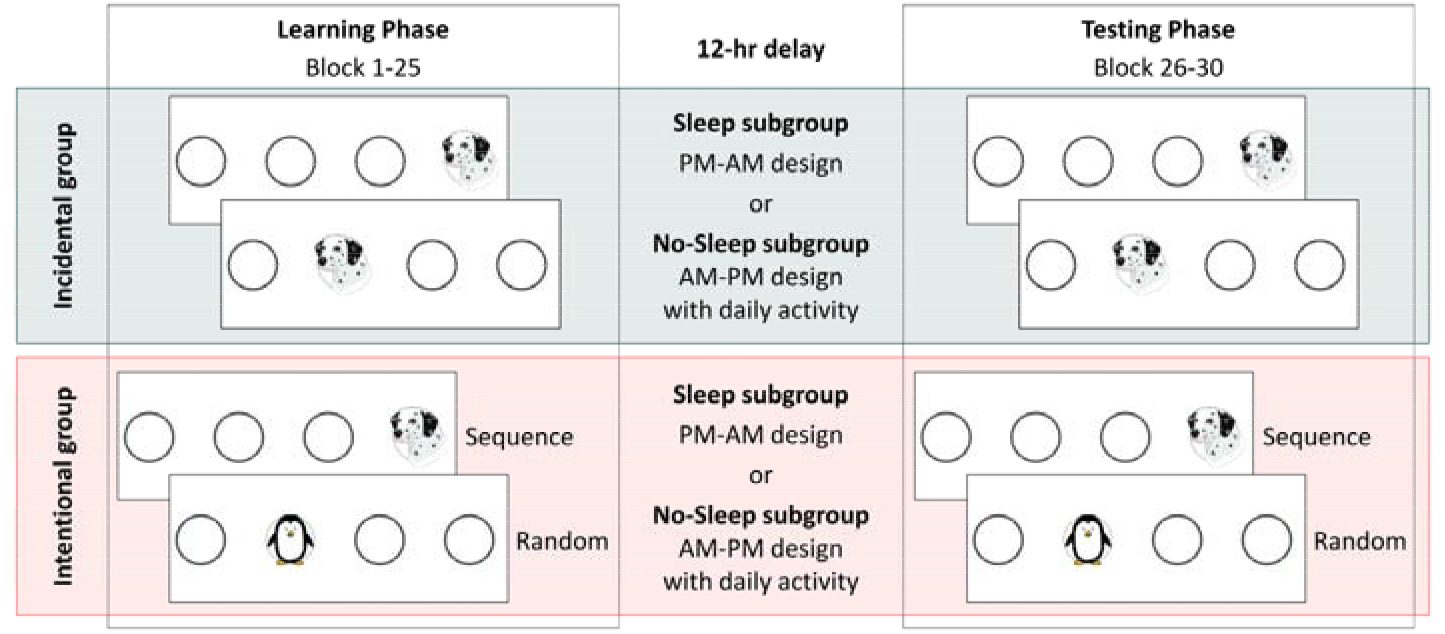
Design and procedure of the experiment. The original fully implicit (Incidental) and the modified cued (Intentional) version of the ASRT task were administered in the experiment. In the cued version of the task (red panel), the regularity was marked by using different stimuli for the sequence elements (a dog’s head) and for the random ones (penguin). In the incidental version of the task (green panel), sequence and random elements were not marked differently (all stimuli were presented by the dog’s head). The Learning Phase (right column) consisted of 25 blocks, while the Testing Phase (left column) contained five blocks. The two sessions were separated by a 12-hour delay. Based on the activity during the delay period, both of the main groups were divided into to a Sleep (PM-AM design) and a No-Sleep subgroup (AM-PM design). All participants performed the same task in the Testing Phase as in the Learning Phase.

### Statistical analysis

Statistical analysis was based on previous studies (J. Howard & Howard, 1997; Kóbor et al., 2018; Nemeth, Janacsek, & Fiser, 2013; Simor et al., 2019) and were carried out by SPSS version 22.0 software (SPSS, IBM). Epochs of five blocks were analyzed instead of single blocks. The Learning Phase consisted of five epochs, while the Testing Phase consisted of one epoch. The Intentional group showed – on average – lower reaction times (RTs) compared to the Incidental group (see Figure S1). To ensure that potential between-group differences in statistical and sequence learning are not due to these general performance differences, we transformed the data by dividing each participants’ raw RT values with their own average performance of the first epoch of the Learning Phase (for a similar approach see: Nitsche et al., 2003). In the next step, we multiplied all data by 100 for easier interpretability and presentation. This way each participants’ performance at the beginning of the Learning Phase was around 100 and changed as the task progressed (see Figure 3, separately for the Incidental and Intentional groups, irrespective to the type of subgroup (Sleep vs. No-sleep)). Note that results concerning raw RTs and accuracy can be found in the Supplementary Results (Table S1, Table S2 respectively). Following the transformation, the Incidental and Intentional groups showed similar RTs (main effect of INSTRUCTION: *F(1, 72)* = 0.012, η*_p_*^2^ = .000, *p* =.911).

**Figure 3.**
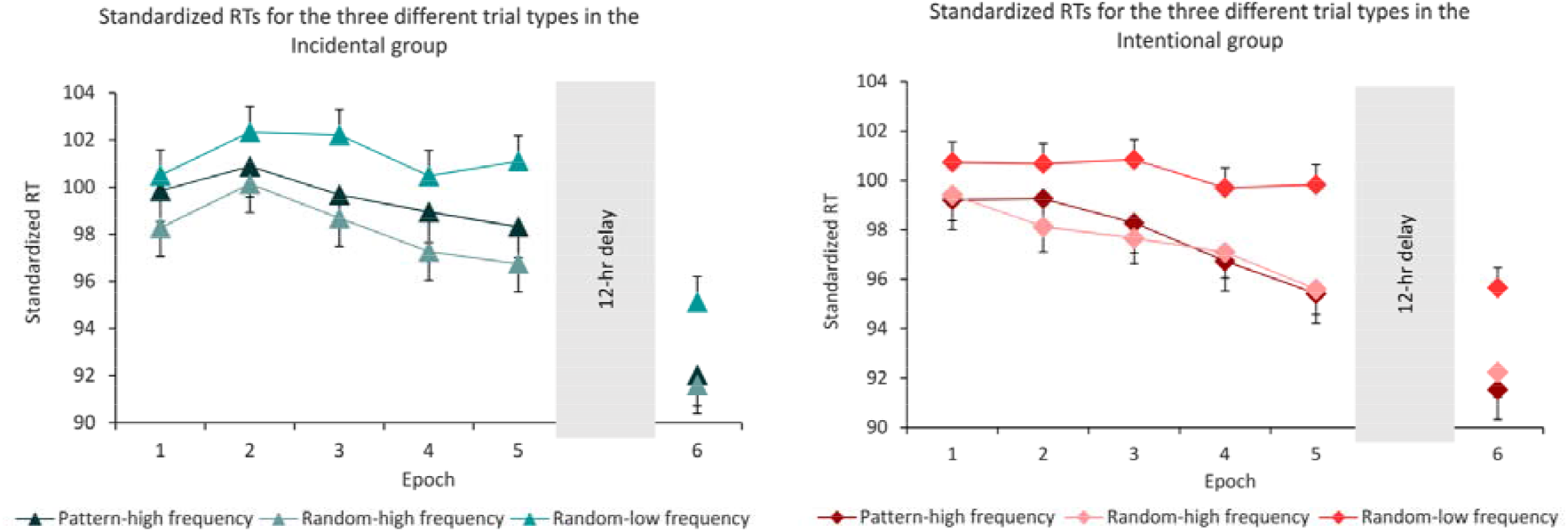
Reaction times (RTs) over the time course of learning for the pattern-high, random-high and random-low frequency elements, separately for the Incidental (green) and the Intentional (red) group, regardless of sleep and no-sleep subgroups. Error bars represent the Standard Error of Mean (SEM).

We calculated median RTs for correct responses only for each participant and each epoch, separately for the pattern, random-high and random-low trial types. Then we calculated learning scores as a difference between RTs (statistical learning: random-low minus random-high, sequence learning: random-high minus pattern). Larger scores indicate better learning performance. Due to the transformation procedure, these learning score can be interpreted as percentages showing how much faster participants responded to the random-high trials compared to the random-low ones (statistical learning) or to the pattern trials compared to the random-high ones (sequence learning). These learning scores were submitted to mixed design ANOVAs to evaluate learning and retention of the acquired knowledge, respectively. Greenhouse-Geisser epsilon (ε) correction was used when necessary. Original *df* values and corrected *p* values (if applicable) are reported together with partial eta-squared (η*_p_*^2^) as the measure of effect size. LSD correction was used for pair-wise comparisons.

As we expected no change in performance over the delay period, except for intentional sequence learning, we conducted Bayesian ANOVAs to overcome the limitations of null-hypothesis significance testing (Dienes, 2011, 2014) and gain statistical evidence for the possible null-results. Bayes Factors (BF) were calculated for the relevant comparisons using JASP (version 0.8.1.1, JASP Team, 2017). In these Bayesian ANOVAs, BF values reflect how well a model behaves compared to the null-model. The smaller the BF_01_ value is, the better the model predicts the data. BF_01_ value of the null-model, which contains the grand mean only is always 1 (Jarosz & Wiley, 2014). Here, we report BF_01_ values for the best fitting model and its ratio to the null-model. The bigger the ratio, the better the alternative model is compared to the null-model.

## Results

### Do the Incidental and Intentional groups show different statistical learning performance in the Learning Phase?

We tested potential group differences in statistical learning between the two groups by conducting a mixed design ANOVA on standardized RT data of the statistical learning scores (i.e., difference between random low-frequency and random high-frequency trial types) with EPOCH (1-5) as a within-subject factor and INSTRUCTION (Incidental vs. Intentional) and SLEEP (Sleep vs. No-sleep) as between-subject factors. Note that for the Learning phase the Sleep vs. No-sleep subgroup comparison can reflect time of day effects rather than the effect of sleep per se, as the Sleep subgroup was first tested in the evening, and the No-sleep subgroup was first tested in the morning. The ANOVA revealed significant statistical learning (INTERCEPT: *F*(1, 70) = 193.807, *p* < .001, η*_p_*^2^ = .735), that is, participants responded faster to random-high frequency trials compared to the random-low frequency ones. The Incidental and Intentional group did not differ significantly in the degree of statistical learning (main effect of INSTRUCTION: *F*(1, 70) = 0.802, *p* = .374, η*_p_*^2^ = .011), indicating that intention to learn did not affect this type of learning. Irrespective of the group instruction, statistical knowledge was increasing as the task progressed (significant main effect of EPOCH: *F*(4, 280) = 5.681, *p* < 0.001, η*_p_*^2^ = .075). The EPOCH x INSTRUCTION interaction did not reach significance (*F*(4, 280) = 0.352, *p* = .839, η*_p_*^2^ = .005), suggesting that the temporal pattern of statistical learning was also independent of intention to learn (see Figure 4).

**Figure 4.**
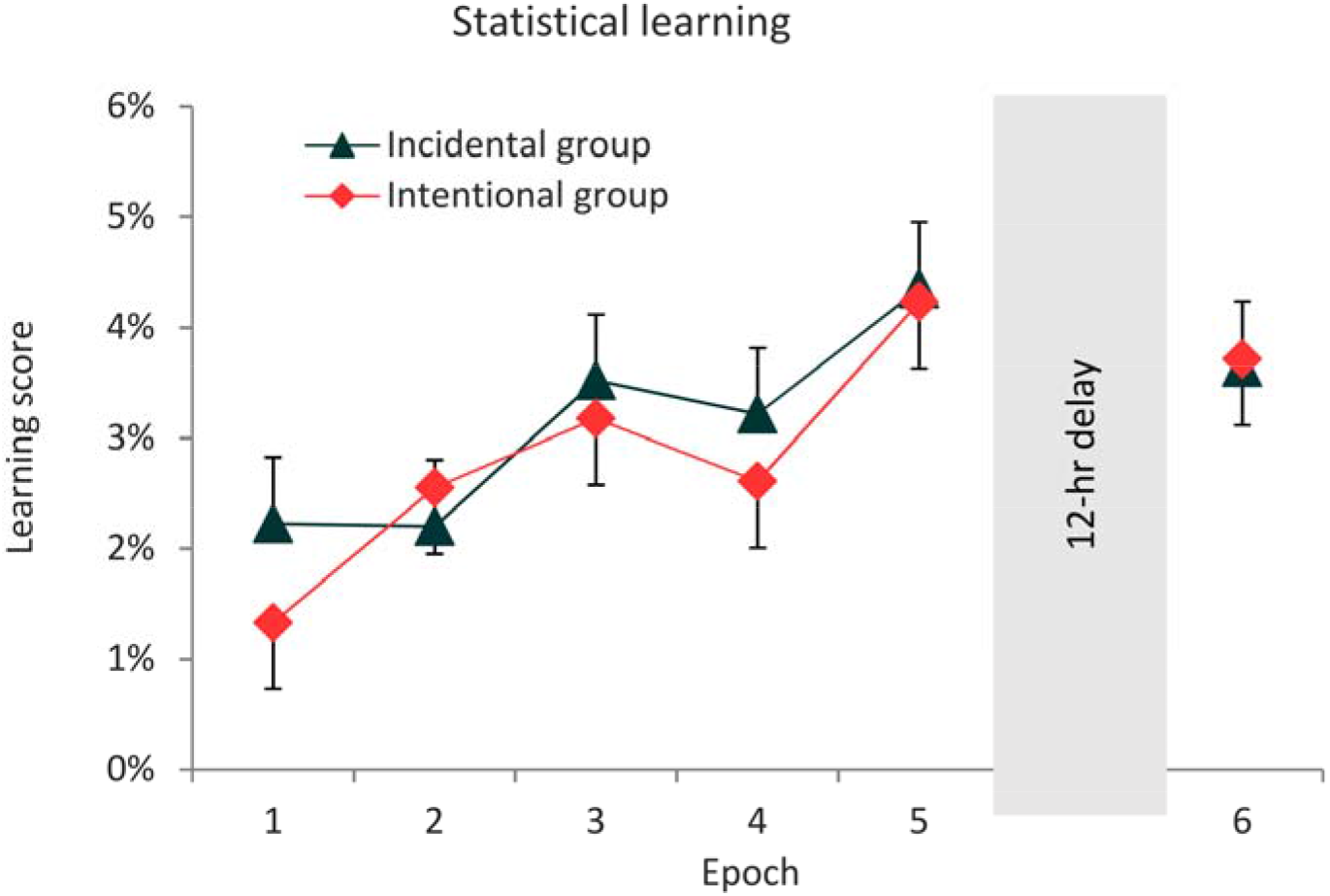
Statistical learning scores (the difference between RTs for random-low vs. random-high frequency trial types) over the Learning and Testing Phases. The Incidental and Intentional groups showed similar learning performance in the first session. Over the 12-hour delay period, both the Incidental and Intentional groups retained the acquired knowledge. Error bars represent the SEM.

The Sleep and No-sleep subgroups, irrespective to the level of intention, showed similar overall degree of statistical learning (main effect of SLEEP: *F*(1, 70) = 0.523, *p* = .472, η*_p_*^2^ = .007) and did not differ in their learning trajectory (EPOCH x SLEEP interaction: *F*(4, 280) = 0.508, *p* = .726, η*_p_*^2^ = .007). However, the INSTRUCTION x SLEEP interaction was significant (*F*(1, 70) = 4.492, *p* = .038, η*_p_*^2^ = .060), indicating that the time of day differently affected incidental and intentional learning performance. Pairwise comparisons revealed that the degree of statistical learning was greater in the Incidental – Sleep group (differing significantly from the Incidental – No-sleep subgroup, *p* = .049, and from Intentional – Sleep subgroup, *p* = .042; all other *p*s > .326; see also Figure S2 in the Supplementary Results). The EPOCH x INSTRUCTION x SLEEP interaction did not reach significance (*F*(4, 280) = 0.161, *p* = .956, η*_p_*^2^ = .002), indicating that the temporal characteristics of statistical learning was comparable in the four subgroups (see the Supplementary Results for Figure S2).

### Do the Incidental and Intentional groups show different statistical performance over the 12-hr offline period?

A similar mixed design ANOVA on statistical learning scores was conducted to assess retention after the offline period with EPOCH (the last epoch of the Learning Phase and the first epoch of the Testing Phase; thus, Epoch 5 vs. Epoch 6) as a within-subject factor and INSTRUCTION (Incidental vs. Intentional) and SLEEP (Sleep vs. No-sleep) as between-subject factors. Overall, participants showed knowledge of the statistical regularities (significant INTERCEPT: *F*(1, 70) = 178.960, *p* < .001, η*_p_*^2^ = .719). Similarly to the Learning Phase, the Incidental and Intentional groups did not differ significantly in the amount of statistical knowledge (main effect of INSTRUCTION: *F*(1, 70) = 0.080, *p* = .779, η*_p_*^2^ = .001). Statistical knowledge was retained over the offline period (main effect of EPOCH: *F*(1, 70) = 2.183, *p* = .144, η*_p_*^2^ = .030), irrespective of the mode of learning (EPOCH x INSTRUCTION interaction: *F*(1, 70) = 0.002, *p* = .968, η*_p_*^2^ < 0.001).

There were no significant differences in overall learning scores between the Sleep and No-sleep subgroups (main effect of SLEEP: *F*(1, 70) = 0.042, *p* = .839, η*_p_*^2^ = .001), irrespective of the mode of learning (INSTRUCTION x SLEEP interaction: *F*(1, 70) = 1.988, *p* = .163, η*_p_*^2^ = .028). Both the Sleep and No-sleep subgroups retained the acquired knowledge over the delay period (EPOCH x SLEEP interaction: *F*(1, 70) = 0.004, *p* = .951, η*_p_*^2^ < .001), similarly in the Incidental and Intentional groups (EPOCH x INSTRUCTION x SLEEP interaction: *F*(1, 70) = 0.492, *p* = .485, η*_p_*^2^ = .007, see Figure S2).

To confirm our interpretations, a Bayesian mixed design ANOVA and BF_01_ values were calculated for learning scores in Epoch 5 and 6. This analysis provided further support for our results: our data favored the null-model (BF_01_ = 1), suggesting the retention of statistical knowledge over the offline period regardless of the delay activity (Sleep vs. No-sleep) and mode of learning (Incidental vs. Intentional; see Figure 4 and Figure S2).

### Do the Incidental and Intentional groups show different sequence learning performance in the Learning Phase?

To measure sequence learning performance in the Learning Phase, we conducted a mixed design ANOVA similar to the one reported above on sequence learning scores (i.e., difference between pattern and. random-high trials) with EPOCH (1-5) as a within-subject factor and INSTRUCTION (Incidental vs. Intentional) and SLEEP (Sleep vs. No-sleep) as between-subject factors. Overall, the ANOVA revealed a significant INTERCEPT (*F*(1, 70) = 18.805, *p* < .001, η*_p_*^2^ = .205), indicating that participants responded differently to pattern and random-high trial types. Moreover, the main effect of INSTRUCTION was also significant (*F*(1, 70) = 10.770, *p* = .002, η*_p_*^2^ = .133): the Intentional group showed a higher degree of sequence learning compared to the Incidental group (see Figure 5). This finding confirms that participants followed the instruction to intentionally learn the sequential regularity in the task. Note that the Incidental group showed a reverse effect (faster RTs for random-high trials), which is typical in incidental ASRT studies (e.g., Nemeth, Janacsek, Király, et al., 2013; Song et al., 2007a). The main effect of EPOCH and the EPOCH x INSTRUCTION interaction were not significant (*F*(4, 280) = 0.300, *p* = .878, η*_p_*^2^ = .004; *F*(4, 280) = 1.404, *p* = .234, η*_p_*^2^ = .020, respectively).

**Figure 5.**
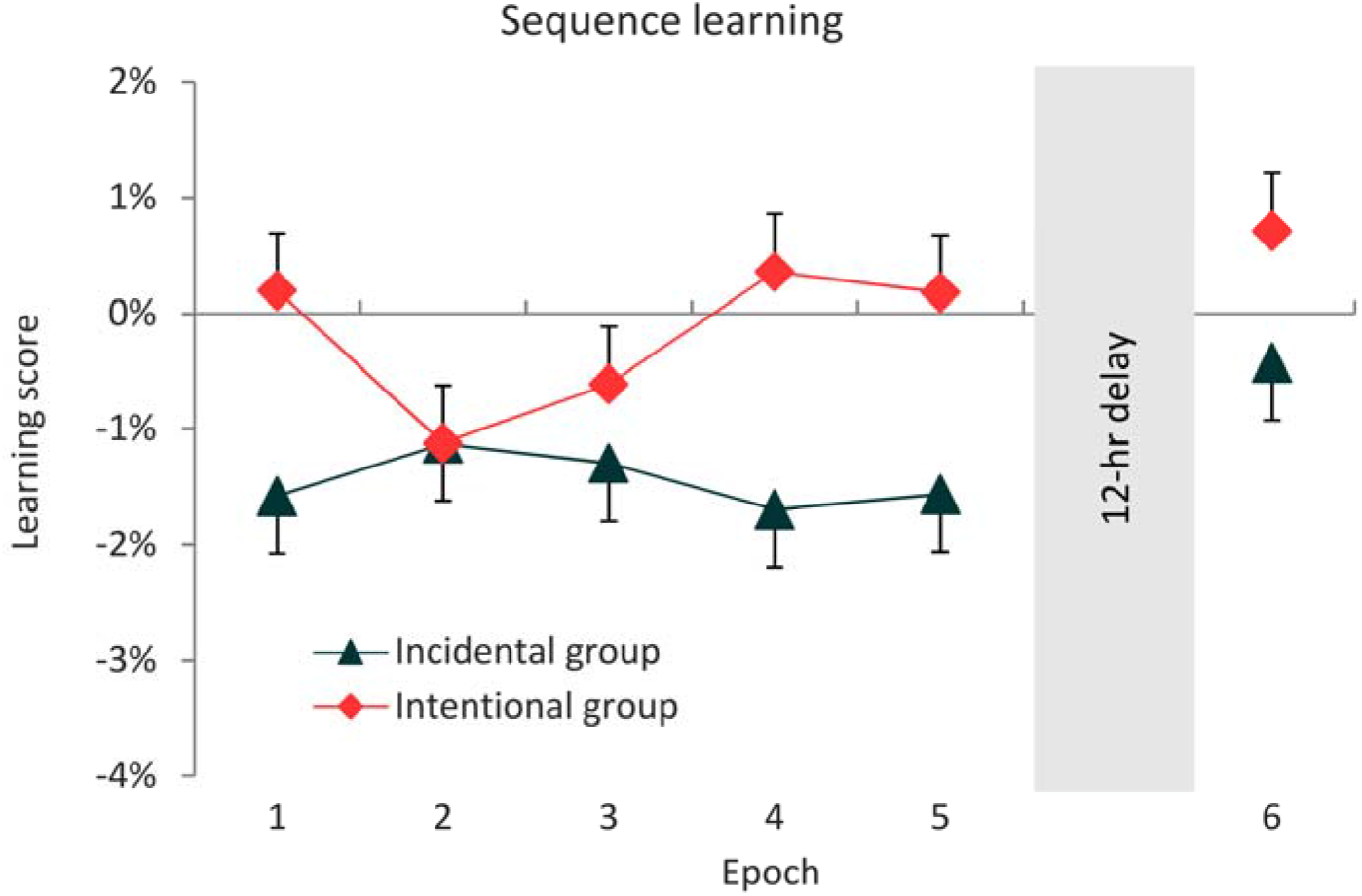
Sequence learning scores (the difference between RTs for pattern vs. random-high frequency trial types) over the Learning and Testing Phases. The Intentional group showed better learning performance in the Learning Phase. Over the 12-hour delay period, both the Incidental and Intentional groups showed no change in the performance. Error bars represent the SEM.

The main effect of SLEEP and the EPOCH x SLEEP interaction did not reach significance (*F*(4, 280) = 3.119, *p* = .082, η*_p_*^2^ = .043, *F*(4, 280) = 0.871, *p* = .479, η*_p_*^2^ = .012, respectively), indicating that the time of day neither affected the overall sequence learning performance, nor its trajectory. The Sleep and No-sleep subgroups within the main groups performed similarly (EPOCH x INSTRUCTION x SLEEP interaction: *F*(4, 280) = 0.242, *p* = .910, η*_p_*^2^ = .003), suggesting that the effect of intention to learn is independent of the time of day (see Figure S3 in the Supplementary Results).

### Do the Incidental and Intentional groups consolidate sequence knowledge differently over the 12-hr offline period?

Offline changes of sequence knowledge over the 12-hr delay were analyzed by comparing sequence learning scores from the last epoch of the Learning Phase and the first epoch of the Testing Phase. These variables were submitted to a mixed design ANOVA with EPOCH (Epoch 5 vs. Epoch 6) as a within-subjects factor and INSTRUCTION (Incidental vs. Intentional) and SLEEP (Sleep vs. No-sleep) as between-subjects factors. The overall sequence learning score did not reach significance (main effect of INTERCEPT: *F*(1, 70) = 0.944, *p* = .335, η*_p_*^2^ = .013). However, the main effect of INSTRUCTION was significant (*F*(1, 70) = 6.966, *p* = .010, η*_p_*^2^ = .091) as the Intentional group showed higher scores compared to the Incidental one, similarly to the Learning Phase. There was a trend for an increase in sequence knowledge over the 12-hour offline period (main effect of EPOCH: *F*(1, 70) = 3.062, *p* = .085, η*_p_*^2^ = .042), similarly in the Incidental and Intentional groups (EPOCH x INSTRUCTION interaction: *F*(1, 70) = 0.428, *p* = .515, η*_p_*^2^ = .006) (see Figure 5).

There were no significant differences in overall learning scores between the Sleep and No-sleep subgroups (main effect of SLEEP: *F*(1, 70) = 0.614, *p* = .436, η*_p_*^2^ = .009), irrespective of the mode of learning (INSTRUCTION x SLEEP interaction: *F*(1, 70) = 0.714, *p* = .401, η*_p_*^2^ = .010). Both the Sleep and No-sleep subgroups retained the acquired knowledge over the delay period (EPOCH x SLEEP interaction: *F*(1, 70) = 0.698, *p* = .406, η*_p_*^2^ = .010), similarly in the Incidental and Intentional groups (EPOCH x INSTRUCTION x SLEEP interaction: *F*(1, 70) = 0.519, *p* = .474, η*_p_*^2^ = .007, see Figure S3).

The Bayesian mixed design ANOVA and BF_01_ values for learning scores in Epoch 5 vs. Epoch 6 revealed that the best fitting model contained the INSTRUCTION factor solely (BF_01_ = 0.31, BF_null-model_ /BF_INSTRUCTION_ = 3.23), the other models had only weaker predictive power. These results provide support for the advantage of the Intentional group in sequence knowledge that seems to be retained over the offline period.

### Was the acquired knowledge intentionally accessible?

#### Sequence reports

On average, participants found and reported the sequence order consequently from the 14^th^ block (*SD* = 7.4). First, we examined the trajectory of consciously accessible sequence knowledge in a repeated measures ANOVA with within-subject factor of EPOCH (1-5). The main effect of EPOCH revealed increasing performance (*F*(4, 136) = 16.90, *p* < .001, η*_p_*^2^ = .332). Pairwise comparison showed that this improvement developed gradually as the task progressed (all *p*s < . 049; *M*_Epoch1_ = 41.8%, *SD*_Epoch1_ = 21.4%, *M*_Epoch5_ = 73.7%, *SD*_Epoch5_ = 31.2%).

Second, we tested whether the sequence report performance changed over the 12-hr offline period by comparing performance on Epoch 5 (last epoch of the Learning Phase) vs. Epoch 6 (Testing Phase) using paired sample t-test. Here we found no significant effect of time (*t*(35) = −1.49, *p* = .146, *d* = 0.096), thus, on the group level, sequence report performance did not change significantly over the 12-hr delay (*M_Epoch 6_* = 76.6%, *SD*_Epoch6_ = 30.3%).

Third, we compared the sequence report performance of the Sleep and No-sleep subgroups to reveal whether the time of day and/or the delay activity affected their intentionally accessible knowledge. Here we found that they performed on a similar level both in the Learning Phase (main effect of SLEEP: *F*(1, 34) = 0.01, *p* = .944, η*_p_*^2^ < .001; SLEEP x EPOCH interaction: *F*(4, 136) = 0.55, *p* = .621, η*_p_*^2^ = .02) and the Testing Phase (both when considering the potential change in percentage of correctly reported elements from Epoch 5 to Epoch 6, SLEEP x EPOCH interaction: *F*(1, 34) = 0.03, *p* = .870, η*_p_*^2^ = .001, and the performance in Epoch 6 separately: *t*(34) = 0.38, *p* = .761, *d* = 0.128).

Finally, we also investigated the relationship of sequence report performance and ASRT task performance. Average sequence report performance significantly correlated with the average sequence learning score in the whole Learning Phase (*r*(35) = .49, *p* = .002), that is better sequence learning performance was associated with better sequence report performance. Similar relationship was found in the Testing Phase (*r*(35) = .35, *p* = .046), thus the positive association remained present after the offline delay. Sequence report performance was not related to statistical learning performance either in the Learning Phase (*r*(35) = .07, *p* = .700) or in the Testing Phase (*r*(35) = .04, *p* = .810).

#### Inclusion/Exclusion task

To test whether the acquired frequency-based information remained implicit or became intentionally accessible, the Inclusion/Exclusion task was administered in both groups (see Methods). Data was analysed in a mixed design ANOVA with CONDITION (Inclusion vs. Exclusion) as a within-subject factor and INSTRUCTION (Incidental vs. Intentional) and SLEEP (Sleep vs. No-sleep) factors as between-subject factors.

The main effect of INTERCEPT (*F*(1, 57) = 65.02, *p* < .0013, η*_p_*^2^ = .533) proved that in overall, participants generated more high-frequency triplets than it would be expected by chance level, which is 25% (Inclusion condition: *M* = 32.2%, *SD* = 0.11%; Exclusion condition: *M* = 31.5%, *SD* = 0.07%). The INSTRUCTION x SLEEP interaction revealed the only other statistically significant effect (*F*(1, 57) = 5.45, *p* = .023, η*_p_*^2^ = .087; all other *p*s > .119). Pairwise comparisons on the *combined* performance in the Inclusion and Exclusion conditions showed that while the Sleep and No-sleep subgroups within the Incidental group performed comparably (*p* = .588), the Intentional – Sleep subgroup generated on average 5.3% more high-frequency triplets compared to the Intentional – No-sleep subgroup (*p* = .009). The lack of significant results concerning the factor of CONDITION indicates that participants generated high-frequency triplets at a similar rate in the Inclusion and Exclusion parts of the task, suggesting that they acquired triplet knowledge, but could not consciously access and control it.

## Discussion

The goal of the present study was twofold. First, we tested how the mode of learning (intentional vs. incidental) affects the subprocesses of procedural memory, namely statistical learning and sequence learning, while controlling for the time on task. Second, we investigated how the post-learning offline period affects the acquired knowledge with a particular focus on the role of the delay activity (sleep vs. wake). Performance was measured by a probabilistic sequence learning task (namely, the ASRT task) and the time on task was controlled by a fixed inter-stimulus interval. Our results revealed that statistical learning was not sensitive to the mode of learning: The Incidental and Intentional groups showed similar learning performance. After the 12-hour post-learning offline period, all groups showed similarly retained statistical knowledge, suggesting that the consolidation was also independent of the mode of learning and was resistant to the delay, irrespective of the delay (sleep vs. wake) activity. In contrast, sequence learning performance was enhanced by intention to learn, and this enhancement remained present after the offline delay as well. The delay activity did not have a differential effect on the consolidation of sequence knowledge. Consolidation effects (i.e., no significant changes in the offline period) were further supported with Bayes Factors. The sequence report task, administered in the Intentional group only, showed that while the ability to consciously recall the sequence order in the ASRT task was associated with better sequence learning performance, there was no such association with statistical learning performance. Finer measures regarding the intentional accessibility of the acquired knowledge (based on the Inclusion-Exclusion task) revealed that both the Incidental and Intentional groups could comparably generate the acquired statistical knowledge, while neither could consciously access and control it.

### Statistical learning and consolidation

The mode of learning had no effect on the *acquisition of statistical knowledge*, in contrast to our hypothesis and the results of Nemeth et al. (2013). The main difference between the two studies is the timing of stimulus presentation. They used self-paced timing, which may have favoured the intentional over the incidental group, as participants in the intentional group could spend more time on task (as much as they needed for a more elaborate stimulus processing). Here we used a fixed ISI to control for the time on task across the intentional and incidental groups. Yet, it is possible that the 500 ms stimulus presentation was too short to exert intentional control over the responses and thus directly influence RTs as opposed to the self-paced version. Future studies seem warranted to examine whether and how various stimulus presentation rates affect statistical learning in intentional vs. incidental learning conditions. As an alternative interpretation, the lack of intention-dependent effect on statistical learning can support the single-system approach of procedural memory (Brown & Robertson, 2007; Cleeremans, 2006; Henke, 2010; Perruchet & Pacton, 2006) and suggests that statistical learning is a uniform process, irrespective of whether learning occurs in an intentional or incidental condition (if time on task is controlled for).

Interestingly, the mode of learning interacted with the *time of day effect*: The Incidental-Sleep subgroup, which completed the Learning Phase during the evening showed somewhat better statistical learning performance compared to the other three groups. So far, only a few studies investigated this question and reported no time of day effects (Durrant et al., 2011; Nemeth et al., 2010). Since this was not the focus of the current study and it is possible that these group differences emerged by chance (as it was not apparent in other measures between these groups), future studies are needed to directly test possible time of day effects on the subprocesses of procedural learning.

Regarding the *consolidation of statistical knowledge*, we found that the acquired information both under intentional and incidental learning conditions was comparably retained during the 12-hour post-learning offline period, irrespective of the delay activity. The consolidation of statistical knowledge has received little empirical attention so far (Durrant, Cairney, & Lewis, 2012; Durrant et al., 2011; Kim et al., 2009). Previous studies that used the incidental version of the ASRT task only focused on the so-called triplet learning measure (see the Task section; Hallgató et al., 2013; Janacsek & Nemeth, 2012; Kóbor et al., 2017; Nemeth & Janacsek, 2010; Nemeth et al., 2010; Song, Howard, & Howard, 2008), which is closely related to statistical learning performance but is somewhat contaminated with sequential information (Nemeth, Janacsek, & Fiser, 2013). The acquired knowledge in these studies seemed to be robustly stable (i.e., neither forgetting, nor improvement occurred) during the offline delay, whether it was short or long, such as 12 hours (Hallgató et al., 2013; Nemeth et al., 2010; Song et al., 2007b), 24 hours (Janacsek, Ambrus, Paulus, Antal, & Nemeth, 2015; Janacsek & Nemeth, 2012), one week (Nemeth & Janacsek, 2010), or even one year (Kóbor et al., 2017; Romano, Howard Jr, & Howard, 2010). Consolidation of (pure) statistical knowledge in the intentional version of the ASRT task has been investigated by Simor et al. (2019), who showed that performance did not change over a 1.5-hour long offline delay. Our findings are consistent with these studies and our hypothesis, showing reliably retained statistical knowledge during a 12-hr offline period, irrespective of whether learning occurred in the intentional or incidental condition.

Sleep did not affect the consolidation of statistical knowledge, as expected based on previous studies (Nemeth et al., 2010; Simor et al., 2019): the acquired knowledge was similarly retained, irrespective of the delay (sleep vs. wake) activity. Contrary, Durrant and colleagues (Durrant et al., 2012, 2011) argue for sleep-dependent consolidation of statistical knowledge, resulting in better performance after a delay period of sleep compared to daytime wakefulness. It is possible that the domain of statistical information has a differential effect on consolidation, as they tested statistical learning in the auditory domain. Altogether, our results show that, at least in the visual domain, statistical knowledge is retained over a 12-hours delay period, irrespective of the mode of learning (intentional vs, incidental) and of the delay activity (sleep vs. wake).

### Sequence learning and consolidation

Regarding *sequence learning*, we found that the instruction to intentionally learn the sequence order together with the additional cued stimulus structure successfully improved performance: The Intentional group reached greater learning scores compared to the Incidental group. This result is consistent with Nemeth et al. (2013), yet, here we measured poorer performance. Despite the instruction and the cued stimuli, the Intentional group only showed near zero-level learning scores, while the Incidental group showed negative learning scores. The former result can be due to the fixed-paced timing. As discussed above, compared to the self-paced timing, where participants could spend more time on task to process the stimuli more elaborately, find the alternating sequence and use that information to improve their performance, the fixed-paced timing may have interfered with one of these processes. The latter finding means that the Incidental group performed better on the high-frequency random trials than on the pattern trials. Previous studies have also reported such result, even using self-paced timing (Nemeth, Janacsek, & Fiser, 2013; Song et al., 2008); the underlying process, however, is still unclear (cf. Kóbor et al., 2019). Better performance on high-frequency random trials may suggest that statistical learning is primary compared to sequence learning. In line with this interpretation, Howard & Howard (2004; 1997) showed that triplet/statistical learning already occurs after one day of practice, while sequence learning develops gradually across a longer practice period (e.g., 4-6 days). Our results altogether show that even though the fixed ISI seems to leed to a decreased sequence learning in the Intentional group compared to previous studies with self-paced timing, the intentional manipulation could still accelerate the process compared to the performance of the Incidental group.

Concerning the *consolidation of sequence knowledge*, we found that the higher performance of the Intentional group remained present during the 12-hour delay as well. Independently of the mode of learning, a slight offline improvement was revealed by a trend in the classical statistical analysis, in line with some of the previous studies (Robertson, Pascual-Leone, & Press, 2004; Spencer et al., 2006). Nonetheless, it was not supported by the Bayesian analysis. Only Simor et al. (2019) investigated the consolidation of sequence knowledge in the intentional, cued version of the ASRT task and found no improvement after the 1.5-hour delay. On the one hand, it is possible that the shorter offline period was not enough to promote improvement of sequence knowledge, while a longer period, such as 12 hours (as in the current study) may be sufficient to reveal small performance improvements. On the other hand, since we found only a trend towards the offline improvement (that was not confirmed by the Bayesian analysis), this result has to be treated with caution and calls for further studies testing under what circumstances this effect may occur.

In contrast to our hypothesis, sleep did not modulate the consolidation of intentionally acquired sequence knowledge. The lack of sleep-effect can support the notion of time-dependent instead of sleep-dependent consolidation in sequence learning (i.e., a simple passage of time is sufficient to consolidate the acquired knowledge, irrespective of the activity during that time). Alternatively, this result may be explained by the fixed-paced timing. As discussed above, it is possible that the 500 ms ISI was too short to intentionally influence performance and the acquired sequence knowledge remained more implicit and less explicit, thus the consolidation effects rather reflect implicit processes as well, which may be less sensitive to the activity in the offline period (Rickard et al., 2008; Robertson, Pascual-Leone, & Press, 2004). This interpretation, however, seems less likely based on the results of the sequence reports (see below).

### Sequence reports

We also examined whether participants could *intentionally recall and control* the acquired sequence knowledge. *Sequence knowledge* of the Intentional group was tested continuously during the entire ASRT task by the *sequence* reports. Considering this measure, we found poorer performance compared to other studies using the intentional, cued version of the ASRT task (Kóbor et al., 2018; Simor et al., 2019), which might also be the result of the temporal parameters of stimulus presentation: Although participants showed a gradually increasing sequence learning performance, the acquired knowledge was not fully available for explicit, consciously accessible processes. Interestingly, sequence report performance was also retained during the offline delay and was not affected by sleep despite the rather explicit nature of the task. Better sequence learning performance on the ASRT task was associated with better sequence report performance, suggesting that although sequence learning was decreased due to the fast stimulus presentation, the acquired knowledge was still intentionally recallable. In the Incidental group, explicit knowledge about the sequence structure was tested at the end of the experiment (in order to keep the learning situation incidental, see Methods). In this group, none of the participants recognized and could intentionally recall the sequence. Altogether, the sequence reports showed that the Intentional group could intentionally recall the order of the sequence and this performance was associated with sequence learning performance in the ASRT task.

### Inclusion-Exclusion task

*Statistical knowledge* was tested by the Inclusion-Exclusion task (Destrebecqz & Cleeremans, 2001; Kóbor et al., 2017) after the Testing Phase in both groups. This measure revealed that participants could generate statistical knowledge, but could not intentionally access and control it, independent of the mode of learning. Interestingly, the Intentional – Sleep subgroup outperformed the other three groups, which may suggest that intention to learn and sleep during the delay period together can benefit the intentional access of statistical knowledge. Overall, this additional measure showed that participants could not intentionally control the knowledge on the frequency-based associations, independently of whether learning occurred incidentally or intentionally.

### Limitations and strengths

Our findings show that the mode of learning (incidental vs. intentional) differentially affects statistical learning and sequence learning processes within procedural memory. A potential limitation of our study may be the temporal parameters of the stimulus presentation. We intentionally chose a fixed ISI to fully control the timing of the presentation, making the intentional and incidental learning conditions comparable in terms of the time spent on the task. Here we conclude that although 500 ms seems sufficient for information processing and responding, this short and fixed ISI still might have been too fast to reach optimal performance. The optimal parameters of the ISI are not clear yet (and may depend on other experimental factors as well), hence further studies are needed to test slower but fixed ISI parameters as well. On the other hand, the present study is uniquely high-powered in the field of comparing intentional/explicit and incidental/implicit learning as we had 37 participants per group compared to the typical group size of 15-20 in the previous studies focusing on learning. Moreover, we involved more participants to investigate the post-learning offline period and the role of delay activity (17-19 participants per subgroups) compared to the sample size of 10-12, which has been typical in previous consolidation studies (e.g., Robertson, Pascual-Leone, & Press, 2004; Walker, Brakefield, Morgan, Hobson, & Stickgold, 2002). Additionally, previous studies mostly used tasks with deterministic sequences, such as in the classical SRT task. It has been shown that the test-retest reliability of the deterministic SRT is weaker (Stark-Inbar, Raza, Taylor, & Ivry, 2016; West, Vadillo, Shanks, & Hulme, 2018), while the probabilistic sequence structure appears to make the ASRT task a more reliable tool to test procedural memory (Stark-Inbar et al., 2016). Altogether, whereas the present study could be improved with more optimal stimulus presentation parameters, the main strengths of the study are the relatively large sample sizes and using a task that showed better reliability compared to other, widely used tasks.

## Conclusions

In summary, our study showed that intention to learn differently affects the parallel formation of two subprocesses of procedural memory, namely statistical learning and sequence learning. Whereas sequence learning can be accelerated with intentional instruction, statistical learning seems to be independent of the mode of learning. However, the consolidation of these two types of information is comparable: both statistical knowledge and sequence knowledge are retained during a 12-hour delay period. Sleep seems to have no superiority compared to a wake delay activity in the consolidation of either statistical or sequence knowledge, suggesting a time-dependent rather than a sleep-dependent consolidation. Overall, our findings provide a deeper insight into how the mode of learning (intentional vs. incidental) and the delay activity in the post-learning period (sleep vs. wake) affects subprocesses of procedural memory formation and consolidation.

## Supporting information

Supplementary results

## Acknowledgement

This research was supported by the National Brain Research Program (2017-1.2.1-NKP-2017-00002 to D.N.); Hungarian Scientific Research Fund (NKFIH OTKA PD 124148 to K.J., NKFIH OTKA K 128016) and Janos Bolyai Research Fellowship of the Hungarian Academy of Sciences (to K.J.). D.N. is helpful for the support of IMÉRA. The authors thank the help of Balázs Török in data acquisition and data analysis.

