## Supplementary results for "Intention to learn differentially affects subprocesses of procedural learning and consolidation: Evidence from a probabilistic sequence learning task"

The Incidental and Intentional groups differed significantly in their average reaction times (RTs) in the Learning phase ($p$ = .051) and over the 12-hour delay ($p$ = .030), see Figure S1. Here we report the ANOVA results for statistical and sequence learning scores calculated on these raw (non-standardized) RT data (see Table S1). Additionally, we present here the ANOVA results for statistical and sequence learning scores calculated on accuracy data (see Table S2). Similar ANOVAs were conducted on the standardized RTs and are presented in the main text of the manuscript. Furthermore, we show here the Sleep and No-Sleep subgroups' statistical learning (Figure S2) and sequence learning performance (Figure S3) calculated on the standardized RT data.

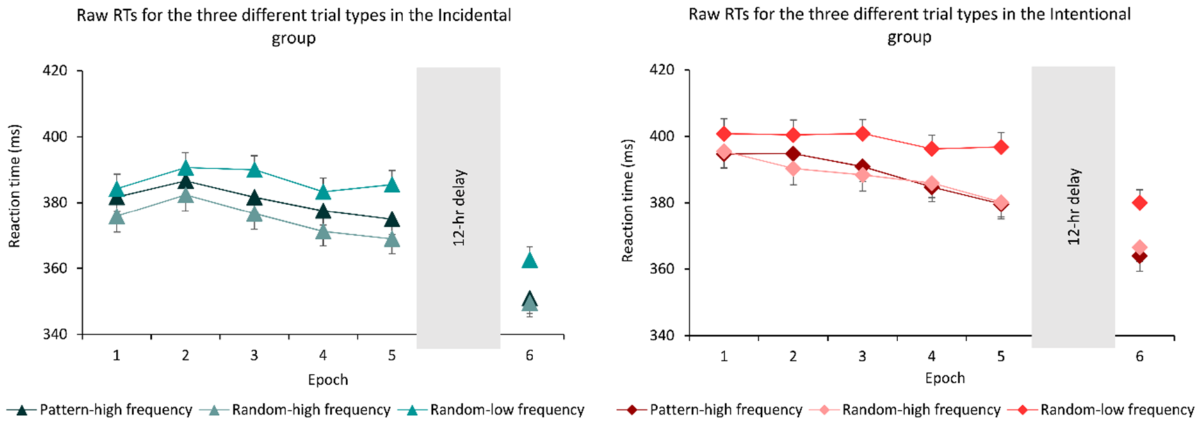


**Figure S1. Raw reaction times (RTs) over the time course of learning** for the pattern-high, random-high and random-low elements, separately for the implicit (greens) and the explicit (reds) group. Error bars represent the Standard Error of Mean (SEM).

**Table S1.** ANOVA results for statistical (A) and sequence (B) learning scores calculated on raw (non-standardized) RT data.

| **A) Statistical learning** | INTERCEPT | | | GROUP | | | EPOCH | | | GROUP x EPOCH | | |
|---|---|---|---|---|---|---|---|---|---|---|---|---|
| | *F* | $\eta^2$ | *p* | *F* | $\eta^2$ | *p* | *F* | $\eta^2$ | *p* | *F* | $\eta^2$ | *p* |
| Learning Phase | 192.663 | .728 | < .001 | 0.249 | .003 | .619 | 5.688 | .073 | < .001 | 0.359 | .005 | .834 |
| 12-hr delay | 178.682 | .713 | < .001 | 0.012 | < .001 | .911 | 2.457 | .033 | .121 | 0.013 | < .001 | .910 |

| | SUBGROUP | | | GROUP x SUBGROUP | | | EPOCH x SUBGROUP | | | GROUP x EPOCH x SUBGROUP | | |
|---|---|---|---|---|---|---|---|---|---|---|---|---|
| | *F* | $\eta^2$ | *p* | *F* | $\eta^2$ | *p* | *F* | $\eta^2$ | *p* | *F* | $\eta^2$ | *p* |
| Learning Phase | 0.371 | .005 | .542 | 4.229 | .057 | .043 | 0.487 | .007 | .741 | 0.192 | .003 | .939 |
| 12-hr delay | 0.002 | < .001 | .964 | 2.095 | .029 | .152 | 0,001 | < .001 | .983 | 0.378 | .005 | .541 |

| **B) Sequence learning** | INTERCEPT | | | GROUP | | | EPOCH | | | GROUP x EPOCH | | |
|---|---|---|---|---|---|---|---|---|---|---|---|---|
| | *F* | $\eta^2$ | *p* | *F* | $\eta^2$ | *p* | *F* | $\eta^2$ | *p* | *F* | $\eta^2$ | *p* |
| Learning Phase | 17.212 | .193 | < .001 | 8.877 | .110 | .004 | .326 | .005 | .853 | 1.332 | .018 | .259 |
| 12-hr delay | 1.014 | .014 | .317 | 6.123 | .078 | .016 | 3.286 | .044 | .074 | 0.489 | .007 | .487 |

| | SUBGROUP | | | GROUP x SUBGROUP | | | EPOCH x SUBGROUP | | | GROUP x EPOCH x SUBGROUP | | |
|---|---|---|---|---|---|---|---|---|---|---|---|---|
| | *F* | $\eta^2$ | *p* | *F* | $\eta^2$ | *p* | *F* | $\eta^2$ | *p* | *F* | $\eta^2$ | *p* |
| Learning Phase | 0.079 | .001 | .779 | 2.723 | .037 | .103 | .797 | .011 | .524 | 0.277 | .004 | .886 |
| 12-hr delay | 0.503 | .007 | .481 | 0.682 | .010 | .412 | 0.642 | .009 | .426 | 0.519 | .007 | .474 |

*Note*. Since the ANOVAs are conducted on learning scores, the INTERCEPT can indicate significant learning. The main effect of GROUP can indicate group differences in the learning scores. The main effect of EPOCH can indicate changes in the learning scores during the task, while the GROUP x EPOCH interaction can indicate group differences in the time course of learning. The main effect of SUBGROUP can indicate Sleep and No-sleep subgroup differences irrespective to the level of intention to learn. The GROUP x SUBGROUP interaction can indicate the subgroup differences within the main groups, while the GROUP x EPOCH x SUBGROUP interaction can indicate these differences in the time course of learning. For more details, see the main text of the manuscript.

**Table S2.** ANOVA results for statistical (A) and sequence (B) learning scores calculated on accuracy data.

| **A) Statistical learning** | INTERCEPT | | | GROUP | | | EPOCH | | | GROUP x EPOCH | | |
|---|---|---|---|---|---|---|---|---|---|---|---|---|
| | *F* | $\eta^2$ | *p* | *F* | $\eta^2$ | *p* | *F* | $\eta^2$ | *p* | *F* | $\eta^2$ | *p* |
| Learning Phase | 62.366 | .464 | < .001 | 1.991 | .027 | .163 | 3.952 | .052 | .005 | 0.887 | .012 | .466 |
| 12-hr delay | 55.444 | .435 | < .001 | 1.619 | .022 | .207 | 0.686 | .009 | .410 | 0.328 | .005 | .568 |

| | SUBGROUP | | | GROUP x SUBGROUP | | | EPOCH x SUBGROUP | | | GROUP x EPOCH x SUBGROUP | | |
|---|---|---|---|---|---|---|---|---|---|---|---|---|
| | *F* | $\eta^2$ | *p* | *F* | $\eta^2$ | *p* | *F* | $\eta^2$ | *p* | *F* | $\eta^2$ | *p* |
| Learning Phase | 1.207 | .017 | .276 | 0.007 | < .001 | .934 | 1.565 | .022 | .188 | 0.643 | .009 | .620 |
| 12-hr delay | 1.268 | .018 | .264 | 0.195 | .003 | .660 | 2.031 | .028 | .159 | 0.544 | .008 | .463 |

| **B) Sequence learning** | INTERCEPT | | | GROUP | | | EPOCH | | | GROUP x EPOCH | | |
|---|---|---|---|---|---|---|---|---|---|---|---|---|
| | *F* | $\eta^2$ | *p* | *F* | $\eta^2$ | *p* | *F* | $\eta^2$ | *p* | *F* | $\eta^2$ | *p* |
| Learning Phase | 2.890 | .039 | .093 | 0.215 | .003 | .644 | 0.616 | .008 | .641 | 1.557 | .021 | .190 |
| 12-hr delay | < 0.001 | < .001 | .986 | 0.450 | .006 | .504 | 1.100 | .020 | .226 | 1.489 | .020 | .226 |

| | SUBGROUP | | | GROUP x SUBGROUP | | | EPOCH x SUBGROUP | | | GROUP x EPOCH x SUBGROUP | | |
|---|---|---|---|---|---|---|---|---|---|---|---|---|
| | *F* | $\eta^2$ | *p* | *F* | $\eta^2$ | *p* | *F* | $\eta^2$ | *p* | *F* | $\eta^2$ | *p* |
| Learning Phase | 0.484 | .007 | .489 | 0.093 | .001 | .761 | 1.229 | .017 | .299 | 0.365 | .005 | .821 |
| 12-hr delay | 0.604 | .009 | .440 | 0.022 | <. .001 | .882 | 3.768 | .051 | .056 | 0.339 | .005 | .562 |

*Note*. For how to interpret the main effects and interaction see the note of Table S1.

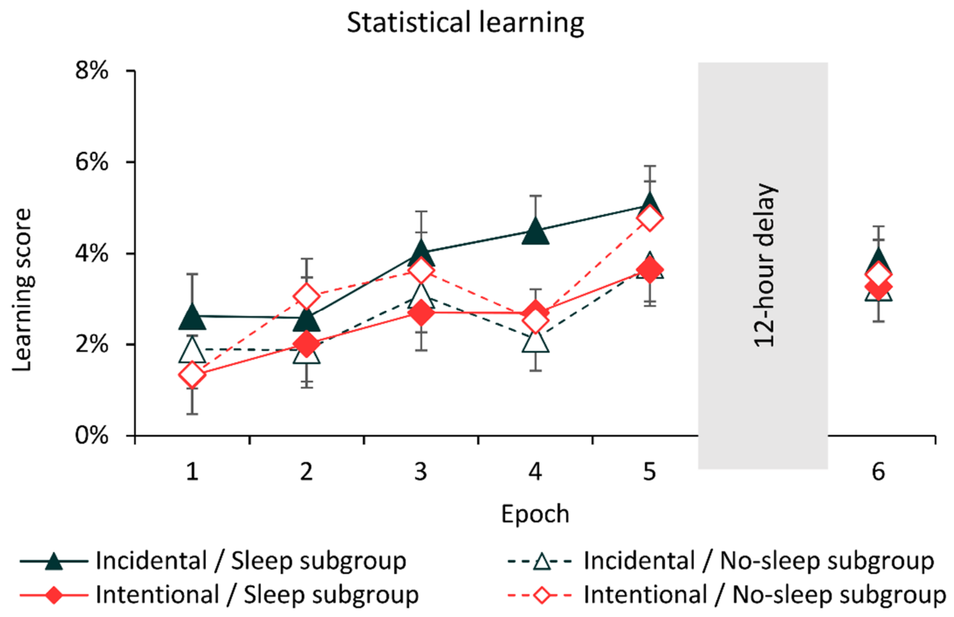


**Figure S2.** Statistical learning scores of the Sleep and No-sleep subgroups over the Learning and Testing Phases (standardized RT data). Error bars represent the SEM.

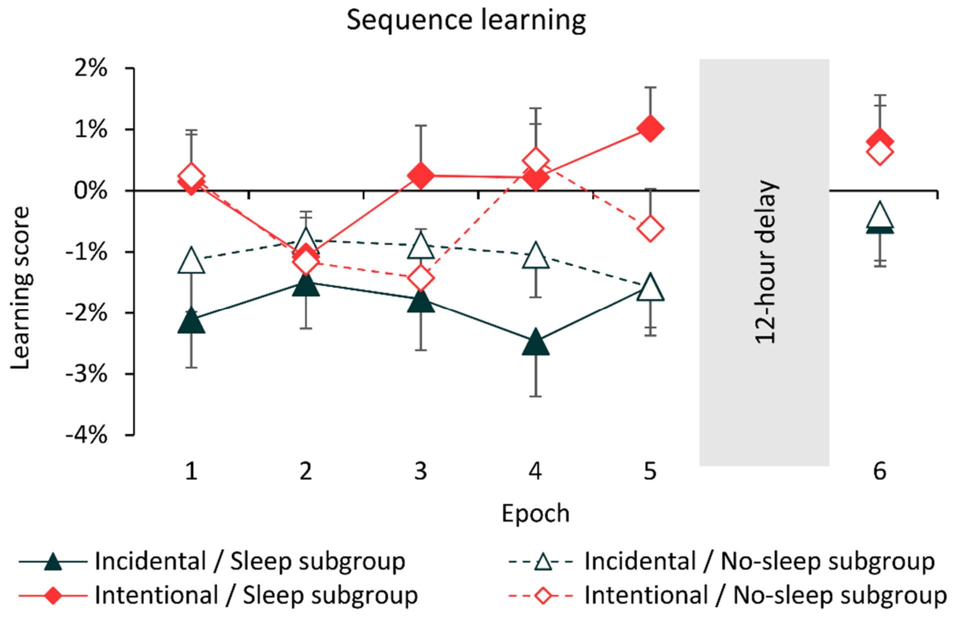


**Figure S3.** Sequence learning scores of the Sleep and No-sleep subgroups over the Learning and Testing Phases (standardized RT data). Error bars represent the SEM.
